# DynaMight: estimating molecular motions with improved reconstruction from cryo-EM images

**DOI:** 10.1101/2023.10.18.562877

**Authors:** Johannes Schwab, Dari Kimanius, Alister Burt, Tom Dendooven, Sjors H.W. Scheres

## Abstract

How to deal with continuously flexing molecules is one of the biggest outstanding challenges in single-particle analysis of proteins from cryo-EM images. Here, we present DynaMight, a new software tool that estimates a continuous space of conformations in a cryo-EM data set by learning 3D deformations of a Gaussian pseudo-atomic model of a consensus structure for every particle image. Inversion of the learnt deformations is then used to obtain an improved reconstruction of the consensus structure. We illustrate the performance of DynaMight for several experimental cryo-EM data sets. We also show how error estimates on the deformations may be obtained by independently training two variational autoencoders (VAEs) on half sets of the cryo-EM data, and how regularisation of the 3D deformations through the use of atomic models may lead to important artefacts due to model bias. DynaMight is distributed as free, open-source software, as part of RELION-5.

---

Structure determination of biological macromolecules by single-particle analysis of electron cryo-microscopy (cryo-EM) images is, at heart, a single-molecule imaging technique. Together, many images of individual complexes in a cryo-EM data set contain information about the full extent of molecular dynamics that existed in the sample when it was plunge frozen. However, stringent low-dose imaging conditions, necessary to limit radiation damage, lead to high levels of experimental noise. Averaging over multiple individual images is thus necessary to extract detailed information about the underlying three-dimensional structures of the macromolecules. Because averaging projection images of distinct structures leads to blurring in the corresponding three-dimensional reconstruction, image classification algorithms are often used to separate cryo-EM data sets into a user-defined number of structurally homogeneous subsets [1]. Despite their effectiveness in handling cryo-EM data sets with a discrete number of conformations, classification algorithms face challenges when continuous molecular motion is present in the sample. Therefore, continuous molecular motions in cryo-EM data sets is often considered a nuisance, rather than a rich source of information about protein dynamics. Manifold embedding [2] represented an early attempt to describe continuous molecular motions in cryo-EM data sets, although application of this approach has been limited to a few macromolecular complexes [3, 4]. A more widely used approach to deal with continuously flexing complexes has been multi-body refinement [5]. Multi-body refinement divides complexes into independently moving rigid bodies through partial signal subtraction [6, 7, 8]. Independent image alignment and reconstruction for each of the individual bodies leads to better maps than a reconstruction of the entire complex that does not take the structural variability into account. However, image alignment requires a minimum size of the individual bodies, which limits the applicability of multi-body refinement to relatively large complexes. More recently, deep convolutional neural networks in the form of variational auto-encoders (VAEs) have been proposed to map projection images into a continuous multi-dimensional latent space [9, 10, 11]. This mapping no longer assumes the presence of a discrete, user-defined number of structures in the data. Moreover, a corresponding decoder network can be used to reconstruct three-dimensional structures for each point in latent space, allowing the creation of movies that describe three-dimensional protein motions by traversing latent space. These approaches have proven useful in exploring continuous molecular motions. However, in contrast to multibody refinement, the majority of them do not lead to improved reconstructed densities for the moving parts.

Two methods have been proposed that aim to analyze continuous molecular motions, while also improving the reconstructed density of the underlying consensus structure. 3D flexible refinement in cryoSPARC uses an auto-decoder to learn deformations that are applied straight to the cryo-EM map [12]. A quasi-Newtonian optimisation algorithm then uses the learnt deformations to improve a reconstruction of the consensus structure. Alternatively, the Zernike3D approach expresses the cryo-EM map in a basis of 3D Zernike polynomials and uses Powell optimisation to find the deformations for each individual particle image [13]. These deformations are then used in a modified algebraic reconstruction technique (ART) algorithm to obtain an improved reconstruction for the consensus structure.

In this study, we present a new approach, coined DynaMight (for ‘exploring protein **Dyna**mics that **Might** improve your map’). Inspired by [10], DynaMight uses Gaussian pseudo-atoms to model the cryo-EM density. The estimation of the conformational variability in the cryo-EM data set is performed by a VAE, where an encoder maps individual cryo-EM images to latent space and a decoder outputs 3D deformations of the Gaussian pseudo-atoms to infer the different conformational states. We introduce a novel decoder architecture that takes the latent vector alongside spatial coordinates as an input and outputs actual displacements. Compared to [10], given a latent representation, the decoder directly represents the function of interest, namely a deformation field. This enables the opportunity to apply priors directly to the deformation field, for which we explore both benefits and pitfalls. A modified filtered backprojection algorithm, that back-projects individual particle images along curves derived from these deformations, then yields an improved density map of the consensus structure.

## Approach

### Description of conformational variability

We describe the *i*th of *N*_*d*_ particle images, *y*_*i*_, with the following forward model:

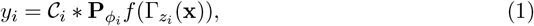

where *𝒞*_*i*_ denotes the contrast transfer function, 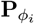 the projection of a particle that is rotated and shifted by its pose *ϕ*_*i*_ *∈ SE*(3). We choose to represent the function *f* by a sum of *N*_*g*_ three-dimensional Gaussian basis functions, or pseudo-atoms:

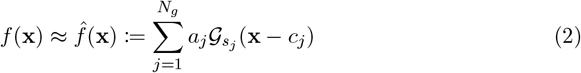

where 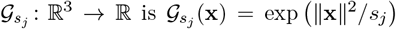. Here *a*_*j*_ *>* 0 denote the amplitudes, *s*_*j*_ *>* 0 the widths, and *c*_*j*_ the central positions of the Gaussian functions.

We assume that all particle images are conformational variations of a single, consensus structure that is described by the *N*_*g*_ three-dimensional Gaussian basis functions, and *z*_*i*_ in equation (1) is the conformational encoding for the *i*th image. We describe the deformation of individual particles as a deviation from the consensus coordinates **x**: Γ(**x**) = **x** *− δ*(**x**), so that:

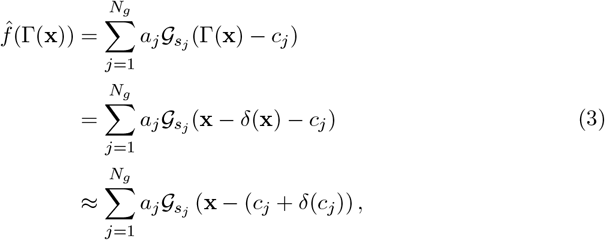

where the last approximation assumes that the deformation field is locally constant and that the density surrounding *c*_*j*_ moves in a similar manner. This enables us to describe the deformations as displacements of the Gaussian centers, which is a computationally tractable representation. Furthermore, the widths *s*_*j*_ and amplitudes *a*_*j*_ of all Gaussian pseudo-atoms are kept the same for the entire dataset. This means that DynaMight is by design constrained to only model mass-conserving heterogeneity and cannot handle non-stoichiometric mixtures. Therefore, compositional heterogeneity should be removed from the data set by alternative approaches, prior to running DynaMight.

### Estimation of conformational variability

For learning the deformations we use a VAE that consist of two neural networks, namely an encoder ℰ that predicts an *l*-dimensional latent representation *z*_*i*_ per particle image, and a decoder 𝒟 that predicts the displacement of all Gaussian pseudo-atoms in the model. The encoder is a fully connected neural network with three linear layers and ReLU activation functions. The input is a (real-space) experimental image *y*_*i*_ and the output are two vectors 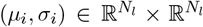, which describe the mean and standard deviation used to generate a sample *z*_*i*_ that serves as input for the decoder.

The decoder 𝒟(*z*_*i*_, *c*_*j*_) then approximates the term *c*_*j*_ + *δ*_*j*_ for each *z*_*i*_. We define the decoder for the entire set of *N*_*g*_ positions as:

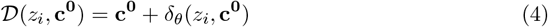

In the above, **c**^**0**^ is all the consensus positions and *δ*_*θ*_ is a differentiable function, 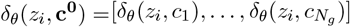, with parameters *θ*, that approximates *δ* for each position. In practice, we evaluate the decoder for each position *c*_*j*_ and query *δ*_*θ*_ with a positional encoding of *c*_*j*_, concatenated with the latent representation *z*_*i*_ that describes the conformation of each particle.

The output positions are used to generate a projection image *p*_*i*_ of the deformed modelin the pose of the particle, and the difference with the experimental image 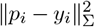 is minimized during training of the neural networks. Once trained, for a latent embedding of the whole data set, one obtains a family of deformation fields 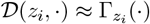 that is defined over the entire three-dimensional space.

### Regularisation and Model Bias

Because of high levels of experimental noise, cryo-EM reconstruction is an ill-posed problem. Even for standard, structurally homogeneous refinement, there are many possible rotational and translational assignments for each image. When estimating conformational variability the poses are known, but many deformed density maps may explain each experimental image equally well. Therefore, in both cases regularisation is essential for robust reconstruction.

The most common form of regularisation in VAEs is to constrain the distribution of latent variables to follow a Gaussian distribution, which lead to the model learning more meaningful and structured representations. The design of the decoder in Figure 1 allows an additional form of regularisation that imposes prior knowledge on its output of real-space deformation fields. A wide range of physically and biologically inspired penalties can be incorporated as priors on the deformations, also see [12, 14, 15]. Possibly a powerful source of prior information would come from an atomic model of the consensus structure, which could provide constraints on chemical bonds, maintain secondary structure elements, etc.

**Figure 1:**
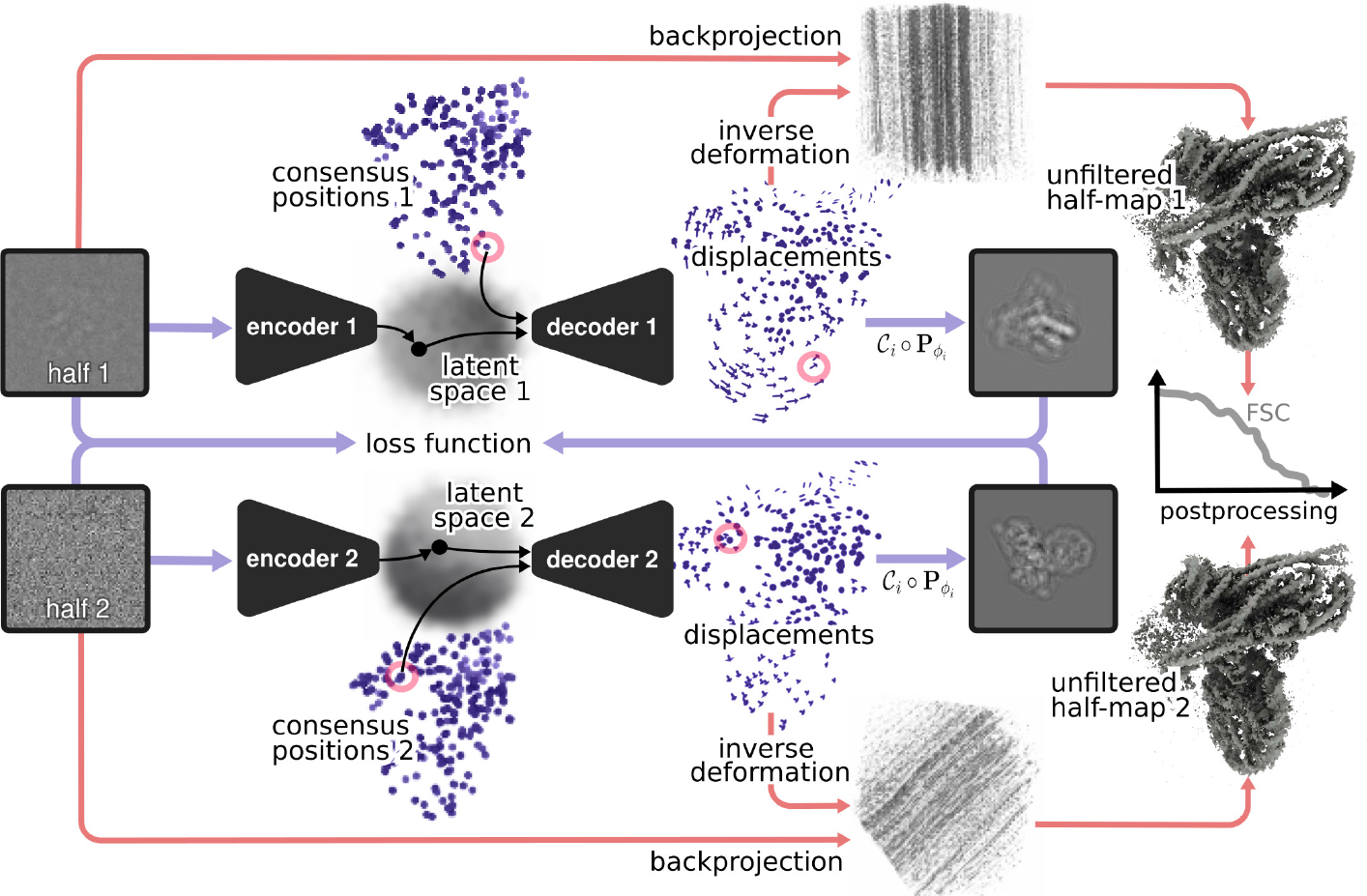
Schematic illustration of DynaMight. Two separate encoders take experimental images from each half set as input, and output a latent vector describing their conformational state. The decoders take the latent vectors together with the coordinates of Gaussian models for the consensus structures for each half set and generate a 3D deformation field for those Gaussians. The deformed models are then projected and compared to the experimental image in the loss function. At the end of the procedure, an approximation to the inverse deformation is used for reconstruction of an improved consensus map for each half set.

To explore direct regularisation of the deformation fields, we tested two approaches. The first approach aims to use prior information from an atomic model that is built in the consensus map, prior to running DynaMight. It generates a coarse-grained Gaussian representation of the atom positions, and then minimizes changes in the distances between these Gaussians according to the bonds that exist in the atomic model:

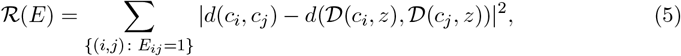

where *E*_*i,j*_ = 1 if there is a bond between the two pseudo-atoms *c*_*i*_ and *c*_*j*_ and *d* denotes Euclidian distance. The deformations with this regularisation scheme result in Gaussians that remain close to a coarse-grained representation of the original atomic model.

The second regularisation approach uses less prior information and does not require an atomic model. Instead, Gaussians are placed to fill densities in the consensus map, and connections *E* in equation 5 are for all pairs of Gaussians that are within a distance of 1.5 times the average distance between all Gaussians and their two nearest neighbours. This regularisation enforces overall smoothness in the deformations. Additional penalties that prevent Gaussians coming too close to each other, or moving too far away from other Gaussians, also exist to ensure a physically plausible distribution of Gaussians.

### Improved 3D reconstruction

We propose an algorithm that uses the estimated deformation fields Γ to obtain an improved reconstruction of the consensus structure that incorporates information from all experimental images. To map back individual particle images to a hypothetical consensus state, one needs to estimate the inverse deformations, which represents a challenge. Whereas the inverse deformation on the displaced Gaussians is given by the negative displacement vector, i.e. Γ^*−*1^(Γ(*c*_*i*_)) = *c*_*i*_, the inverse deformation field needs to be inferred at all Cartesian grid positions of the improved reconstruction. We train a neural network as a regression function to estimate a deformation field that coincides on the given sampling points Γ(*c*_*i*_), but can be evaluated on arbitrary positions. This network consists of an MLP with 6 layers and a single additive residual connection to the original coordinates of the consensus model **c**^**0**^. Similar to the forward deformation model, the network takes the latent code *z*_*i*_ and the deformed positions Γ(*c*_*i*_) as inputs and aims to output the original positions *c*_*i*_. In addition to the inversion of the forward fields on the sampling points, we force the inverse field to be smooth by adding a regularisation term to the loss function.

The algorithm aims to improve the reconstruction of the density *f*, using the known deformations Γ, i.e. we aim to find the minimizer 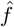 of the data fidelity

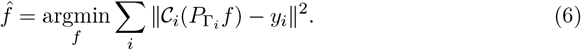

This minimizer can be computed using the reconstruction formula

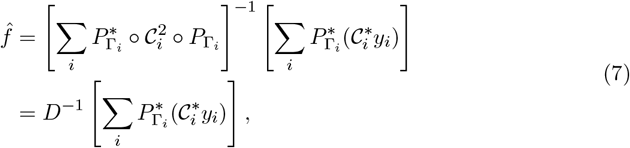

to get an estimate of the unknown density *f*. Here *D* is a matrix that depends on the estimated deformations, and 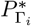 is the composition of the backprojection operator and the inverse deformation corresponding to the *i*th particle. (Figure 1). For the structurally homogeneous case, Γ is the identity operator and *D* is diagonal in Fourier space and therefore the inverse can be computed simply by division, given that the distribution of projection directions covers the whole frequency domain and *D* has no zeros in the diagonal. In the presence of deformations, this matrix is not diagonal anymore and would be too expensive to compute or store. We approximate equation (7) by using the filter that would correspond to the homogeneous case. Although even in the optimal scenario of having complete data of clean projection images, this method does not yield a minimum of functional (6), it still allows to correct for the deformation to some degree. When the deformation fields are not smooth, for example when two nearby domains move in opposite directions, reconstruction with the proposed algorithm may introduce artifacts at the interface between the domains.

### Implementation details

The initial positions of the Gaussians for the VAE are obtained by approximating a map from a consensus refinement with a Gaussian model. This initial consensus map does not correspond to an actual state of the complex, but rather to a mixture of different conformations. Therefore, parts of the map will have regions of poorly defined density, and correspondingly fewer Gaussians. To overcome this limitation, we update the positions of the consensus Gaussian model throughout the estimation of the deformations, such that the positions *c*_*j*_ may correspond to a single conformation at the end of the iterative process.

After initialisation of the Gaussians, in the first epochs of the training of the VAE, we only optimize the global Gaussian parameters, i.e. their widths, amplitudes and positions. These parameters are optimized with the ADAM optimizer and a learning rate of 0.0001. After this initial warm-up phase we start optimisation of the network parameters of the VAEs, again using the ADAM optimizer with a learning rate of 0.0001. During the second phase, the parameters of the Gaussians continue to be updated. Training of the VAEs is stopped when the updates of the consensus model don’t yield improvements anymore or a fixed, userdefined number of epochs are completed.

Training of the VAE is performed on two half sets, where two encoder-decoder pairs are trained independently, as illustrated in Figure 1. This procedure yields two independent families of deformation fields, one for each half-set. The approximate inverse of these deformations are then used using the deformed weighted backprojection algorithm to generate two independent maps with improved estimates for the consensus structure. These half-maps can than be used in conventional post-processing and resolution estimation routines. As described in the Discussion section, by setting aside a small validation set of images, the two independent decoders also allow an error estimation of the displacement fields.

DynaMight has been implemented in pyTorch [16], and is accessible as a separate jobtype from the RELION-5 GUI. DynaMight uses the Napari viewer [17] to visualise the distribution of particles in latent space, as well as the corresponding deformation fields. The same viewer also allows real-time generation of densities from points in latent space, movie generation, and the selection of particle subsets in latent space.

## Results

### Regularisation of the deformations is necessary but can lead to model bias

We first analyzed the different options for regularisation of the deformations on a well characterised data set on the yeast *Saccharomyces cerevisiae* pre-catalytic B complex spliceosome [18](EMPIAR-10180, [19]). The same data, or subsets of it, have also been analyzed using multi-body refinement [5] cryoDRGN [9], Zernike3D [13] and [10]. To minimize computational costs and to ensure structural homogeneity [9], we used 3D classification in RELION [20] to select ∼ 45*k* particles with reasonable density for the head region. Training of the VAEs on this subset with a box size of 320 took about 2.5 minutes per epoch on a single NVIDIA A100 GPU. This resulted in training times between 8 and 12 hours for estimating the deformations. Further estimation of the inverse deformations took ∼ 4 hours and reconstruction with the deformed backprojection ∼ 3 hours on the same GPU.

Without any regularisation of the deformations, estimated deformation fields displayed rapidly changing directions for neighbouring Gaussians, and deformed backprojection yielded reconstructions for which the local resolution did not improve with respect to the original consensus reconstruction (*cf* Figs 2A-B). A consensus reconstruction with better local resolutions was obtained using the regularisation scheme that enforces smoothness in the deformations, but without using an atomic model (Fig 2C). The map with the highest local resolutions was obtained using the regularisation scheme that enforces distances between bonded atoms of an atomic model (PDB-ID: 5nrl) (Fig 2D). It thus appeared that incorporation of prior knowledge from the atomic model into the VAE had been beneficial.

**Figure 2:**
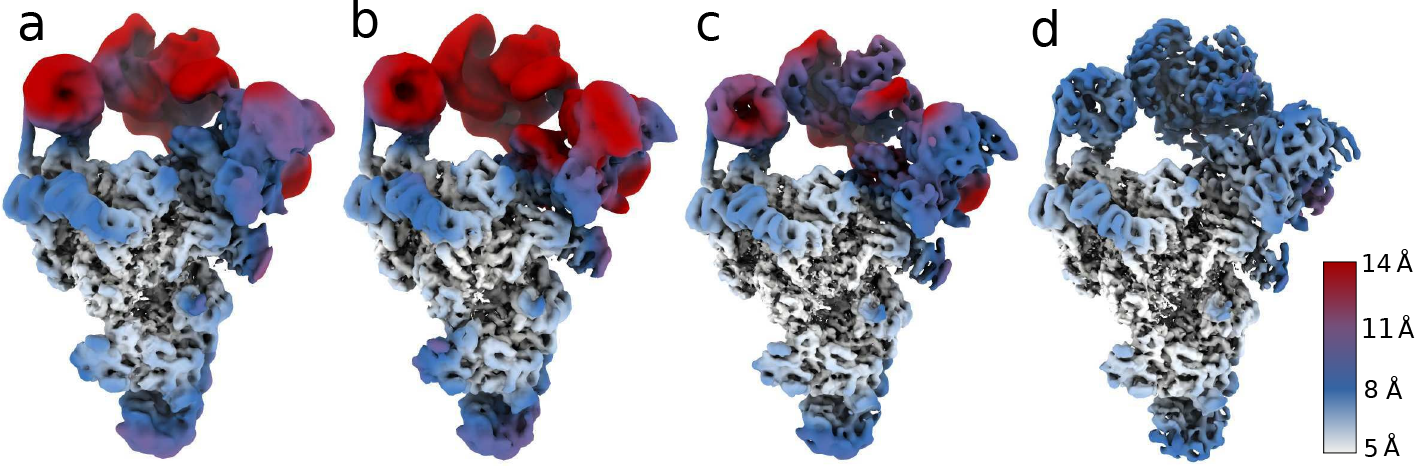
DynaMight reconstructions of the spliceosome subset. **a** Standard RELION consensus refinement; **b** DynaMight without regularisation; **c** DynaMight with smoothness regularisation on the Gaussians; **d** DynaMight with regularisation from an atomic model. All maps are coloured according to local resolution, as indicated by the colour bar.

However, because the neural networks in our approach comprise many parameters, we were worried that there would be scope for “Einstein-from-noise” artefacts, similar to those described for orientational assignments in single-particle analysis [21, 22, 23]. To test this, we performed two control experiments.

In the first control experiment, we replaced the atomic model of the U2 3’ domain/SF3a domain with a different protein domain of similar size (PDB-ID 7YUY) [24]). The U2 3’ domain/SF3a showed only weak density in the consensus map, indicating large amounts of structural heterogeneity in this region. Although using the incorrect atomic model to estimate the deformation fields led to a similar improvement in local resolution compared to using the correct model (*cf* Figs 3A-B), the reconstructed density from the deformed backprojection resembled the incorrect model, rather than the correct model (Fig 3C).

**Figure 3:**
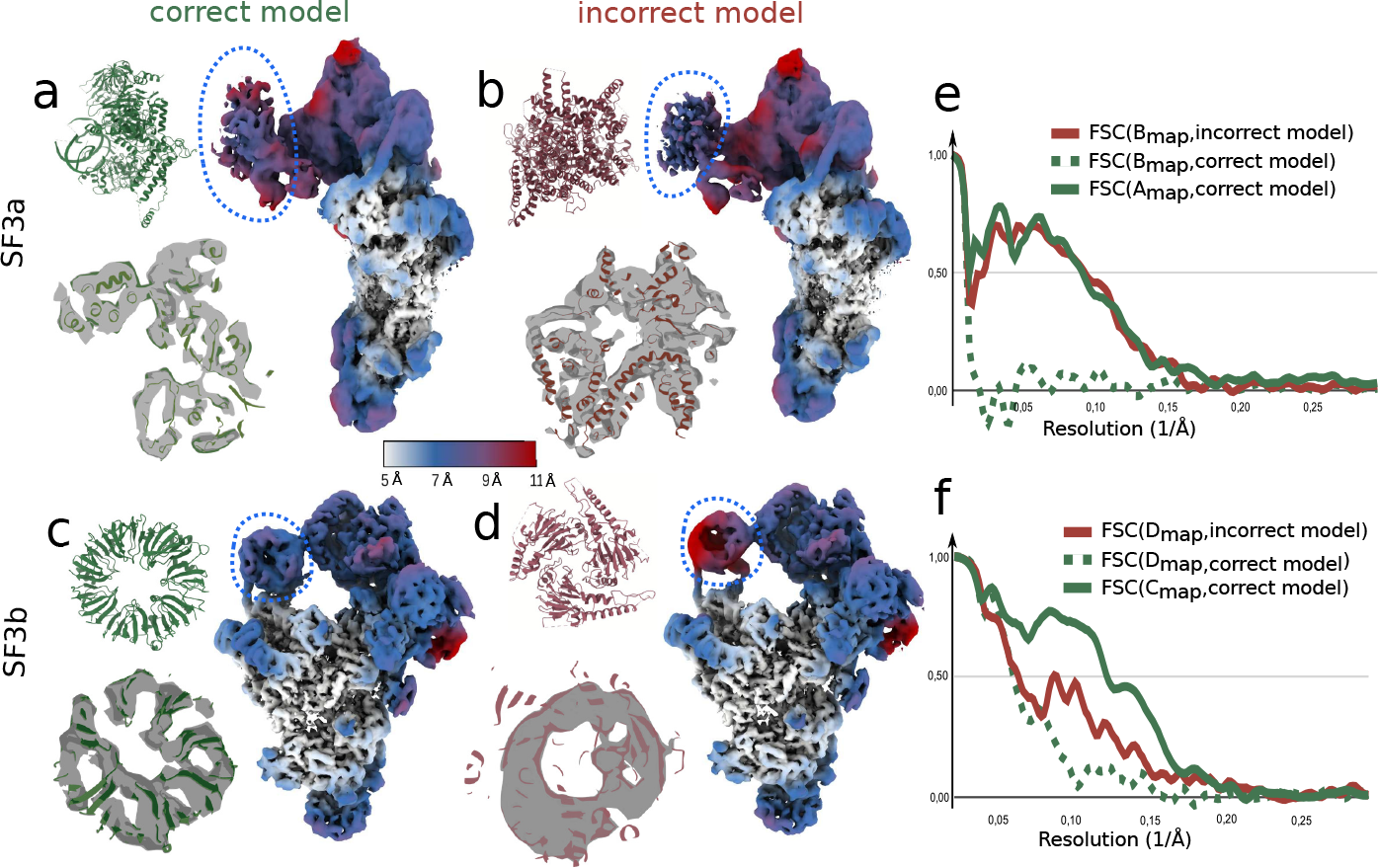
Using incorrect atomic models in DynaMight. **A** Reconstruction after deformed backprojection using the correct atomic model for the SF3a region, colored by local resolution (right). The correct atomic model for the SF3a region is shown in green in the top left; an overlay of that model with the reconstructed density after deformed backprojection is shown in the bottom left. **B**: As in A, but using an incorrect atomic model for the SF3a region (shown in red). **C**: FSC curves between the maps in panels A or B, masked around the SF3a region, and the correct (green) or incorrect (red) atomic models. **D-F** As in A-C, but using different atomic models for the SF3b region.

In the second control experiment, we replaced the atomic model of the SF3b domain with PDB-ID 1G88 [25]. The density for the SF3b domain in the consensus map was stronger than the density for the SF3a region, indicating that this region in the spliceosome is less flexible. In this case, using the incorrect atomic model yielded a map with lower local resolutions in the SF3b region than using the correct model (*cf* Figs 3D-E). But still, the reconstructed density from the deformed backprojection resembled the incorrect model more than the correct model (Fig 3F).

These results suggest that estimation of deformation fields may lead to model bias, to the extent that reconstructed density may reproduce features of an incorrect atomic model. The scope for model bias to affect the deformed backprojection reconstruction is larger in regions of the map with higher levels of structural heterogeneity. Because it would be difficult to distinguish correct atomic models from incorrect ones, we caution against the use of this type of regularisation in DynaMight. Therefore, in what follows, we only used the less informative, smoothness prior on the deformations.

### DynaMight estimates motion and improves maps of inner kinetochore complexes

Next, we demonstrate the usefulness of DynaMight on two cryo-EM data sets of the yeast inner kinetochore [26].

The first data set comprises 100,311 particles of the monomeric constitutive centromere associated network (CCAN) complex bound to a CENP-A nucleosome. For this data set, we trained the half-set VAEs for 220 epochs and we used a 10-dimensional latent space. The estimated 3D deformations are distributed uniformly in latent space (Fig 4A), without specifically clustered conformational states, suggesting that the motions in the data set are mainly of a continuous nature. Analysis of the motions revealed that the nucleosome is rotating in different directions relative to the rest of the complex, and that these rotations co-exist with the up and down bending of the Nkp1, Nkp2, CENP-Q and CENP-U subunits (arrows in Figure 4B and Supplementary Video 1). The reconstruction from deformed backprojection improved local resolutions compared to the consensus map from standard RELION refinement, with clear improvements in the features for both protein and DNA (Fig 4C).

**Figure 4:**
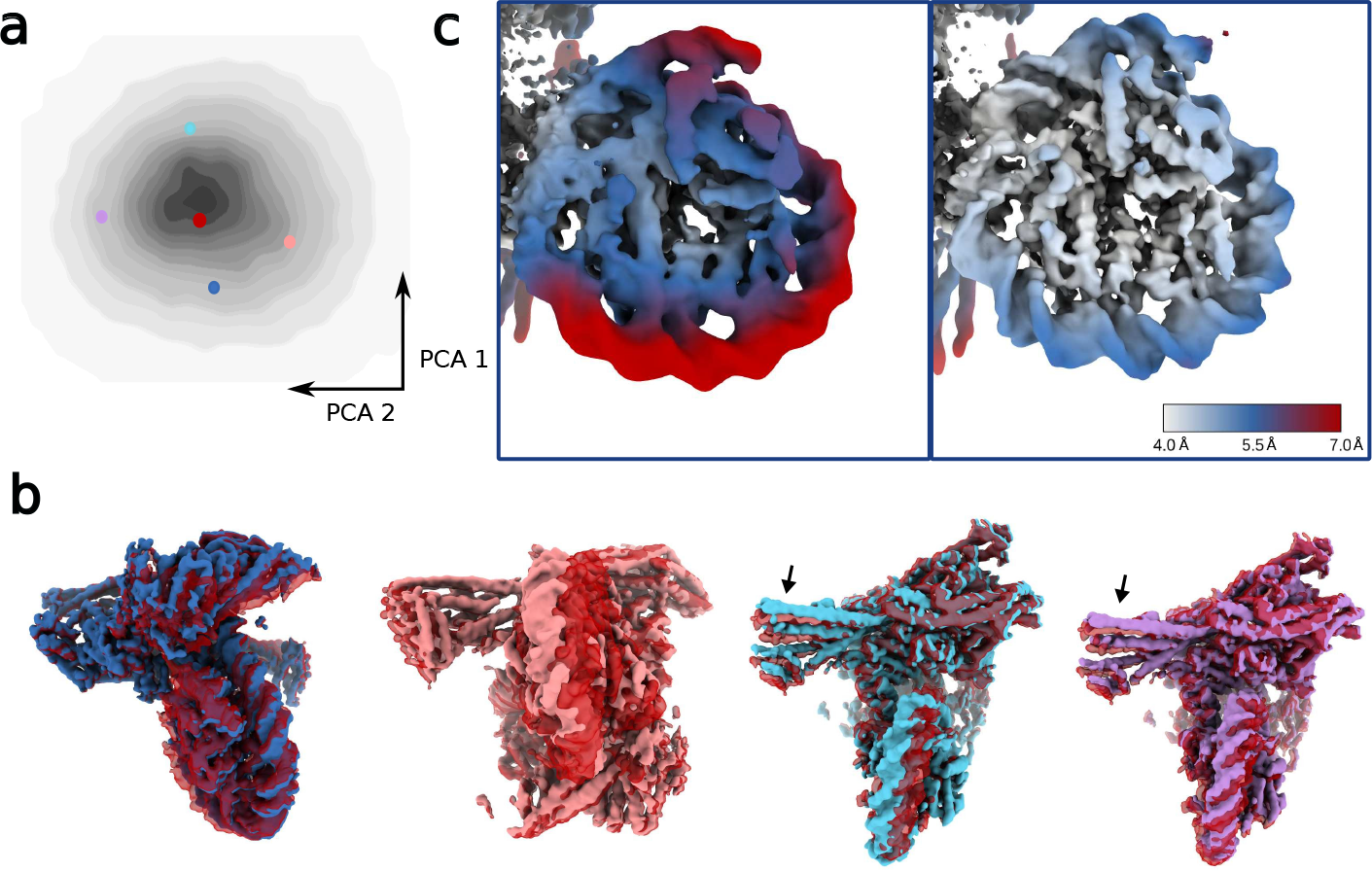
DynaMight results for the monomeric CCAN complex. **a:** PCA of the conformational latent space, with coloured dots indicating the positions of the five maps in panel b. **b** Five conformational states of the complex. One state, in red, is shown in all four panels. The colours of the five maps are the same as the colours of their corresponding dots in panel a. The **c:** Reconstructions from standard RELION consensus refinement (left) and the improved reconstruction using DynaMight (right). Both maps are coloured according to local resolution, as indicated by the colour bar.

The second data comprises 108,672 particles of the complete yeast inner kinetochore complex assembled onto the CENP-A nucleosome. Training of the VAE was done for 290 epochs, and the dimensionality of the latent space was again set to 10. Again, a continuous distribution of deformations in latent space suggests continuous structural flexibility (Fig 5A). Analysis of the deformations revealed large relative motions between different regions of the complex. Different states of the complex are depicted in Fig 5A and Supplementary Video 2. Deformed backprojection resulted in a map with improved local resolution and protein and DNA features compared to the map from consensus refinement (*cf*. Fig 5B-C).

**Figure 5:**
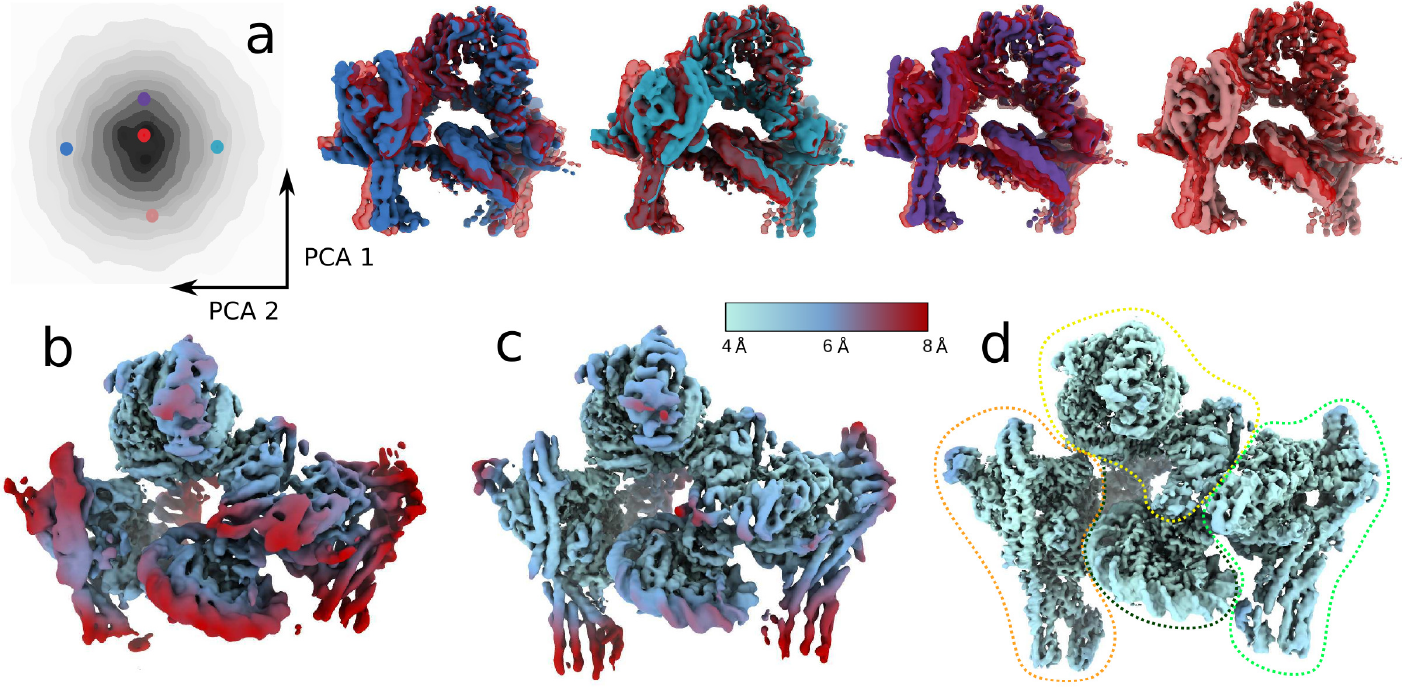
DynaMight results for the complete kinetochore complex. **a:** PCA of the conformational space (on the left) with highlighted positions of five conformation states, the maps of which are shown in the same colours on the right. **b:** Maps from standard RELION consensus refinement, **c:** DynaMight reconstruction, and **d:** reconstruction using RELION multibody refinement. The outline regions in the latter show the four bodies that were used for multibody refinement. The maps in **b-d** are colored by local resolution, as indicated by the colour bar.

Because this complex, with a molecular weight of 1.5 MDa, is large enough to divide into multiple independently moving rigid bodies, we also applied multi-body refinement [5] to this data set. We used the four bodies illustrated in Fig 5D; body 1 (orange): CCAN^Topo^, body 2 (light green):, 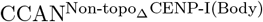, body 3 (yellow): CBF3^Core^+CENP-I^Body^ and body 4 (dark green): CENP-A^Nuc^). The local resolutions resulting from multi-body refinement (Fig 5D) are better than those from the deformed backprojection reconstruction of DynaMight, illustrating that there is still room for further development of the latter.

Training of the VAEs took around 17 and 27 hours on an NVIDIA A100 GPU for the monomeric and dimeric datasets with particle box sizes of 320 and 360. Estimating the inverse deformations took ∼ 6 hours and performing the reconstruction ∼ 9 and ∼ 13 hours respectively.

## Discussion

How to deal with continuous conformational heterogeneity remains a rapidly developing topic in cryo-EM single-particle analysis. As outlined in the Introduction, and recently reviewed in [27], multiple approaches from different labs have been proposed. In this paper we present a new approach, called DynaMight, which consists of two VAEs that are trained independently on half-sets to estimate displacements of a Gaussian model and a modified weighted backprojection algorithm to correct for the estimated deformations. We show for two data sets on the yeast inner kinetochore that DynaMight is useful in improving cryo-EM maps of macromolecular complexes that exhibit large amounts of flexibility. However, as we also show in this paper, there is still obvious scope for further improvements, of DynaMight in particular, and how to deal with continuous structural heterogeneity in general.

Because of the high levels of experimental noise and the large amount of parameters needed to describe continuous structural flexibility in the particles, an obvious way to improve these methods is the incorporation of prior knowledge. However, our results on the spliceosomal B complex show that such approaches are not without risk. We observe that there are enough parameters in DynaMight’s neural networks to result in deformation fields that, when used in deformed backprojection, will reproduce incorrect features from the consensus model that is used to regularise these deformations. That model bias may play a role is perhaps not surprising, given that similar observations have been made for standard (structurally homogeneous) refinement, where only five parameters (three rotations and two translations) are used for every particle. The total number of parameters in DynaMight’s VAE is approximately 10 million, which results in considerably higher numbers of parameter per particle for typical data sets. We do not believe that the risk of overfitting exists only in DynaMight. Other approaches that describe structural heterogeneity in the data set with large neural networks, or other approaches with high numbers of parameters per particle, like cryoDRGN [9], Zernike3D [13], Flex3D [12], will probably also be susceptible to these problems. The development of validation procedures will thus be important. In DynaMight, we chose to not use atomic models for regularisation of the deformations, as potential model bias towards those models takes away the possibility to validate the map by the appearance of protein-like features. The exploration of more sophisticated methods, where part of the information of atomic models is used and other parts are set aside for validation, may yield better methods, while still allowing proper validation.

Because model bias may affect the estimation of deformation fields, over-estimation of the resolution of reconstructions that correct for these deformations may represent another pitfall. Resolutions are typically measured by Fourier shell correlation between two half-sets. However, if deformations have been estimated jointly for both half-sets, with the same reference map as origin, then incorrect features from the reference model may be reproduced in both half-reconstructions, resulting in inflated FSC curves and over-estimation of resolution. By training two independent VAEs with separate consensus models for both half-sets, similar to “gold-standard” approaches in standard refinement [28, 29], this risk is avoided in DynaMight.

Training two VAEs independently on two half-sets of the data also offers an opportunity to estimate the uncertainty in the estimated deformations. Although in recent years multiple methods have been proposed to analyse molecular motions in cryo-EM data sets, less consideration has been given to what extent these motions can be trusted. Error estimates on the deformations can be obtained for a subset of the particles (we used 10% in Figure 6), by excluding this subset from the training of the decoders, and only using it for training its embedding to latent space. For each particle in this subset, one obtains an embedding with both separate encoders to obtain a latent representation for the corresponding decoder. Applying both decoders to get the displacements of either of the consensus models then leads to two independent estimates of the deformations for the particles in the subset. The difference between these two estimates provide an estimate of the errors in them. We illustrate this procedure in Fig 6B, where we observe that the errors in the deformations vary among particles and among different regions of the monomeric CCAN-CENP-A nucleosome complex. Future developments in regularisation methods as described above may benefit from considering estimated errors in the deformations.

**Figure 6:**
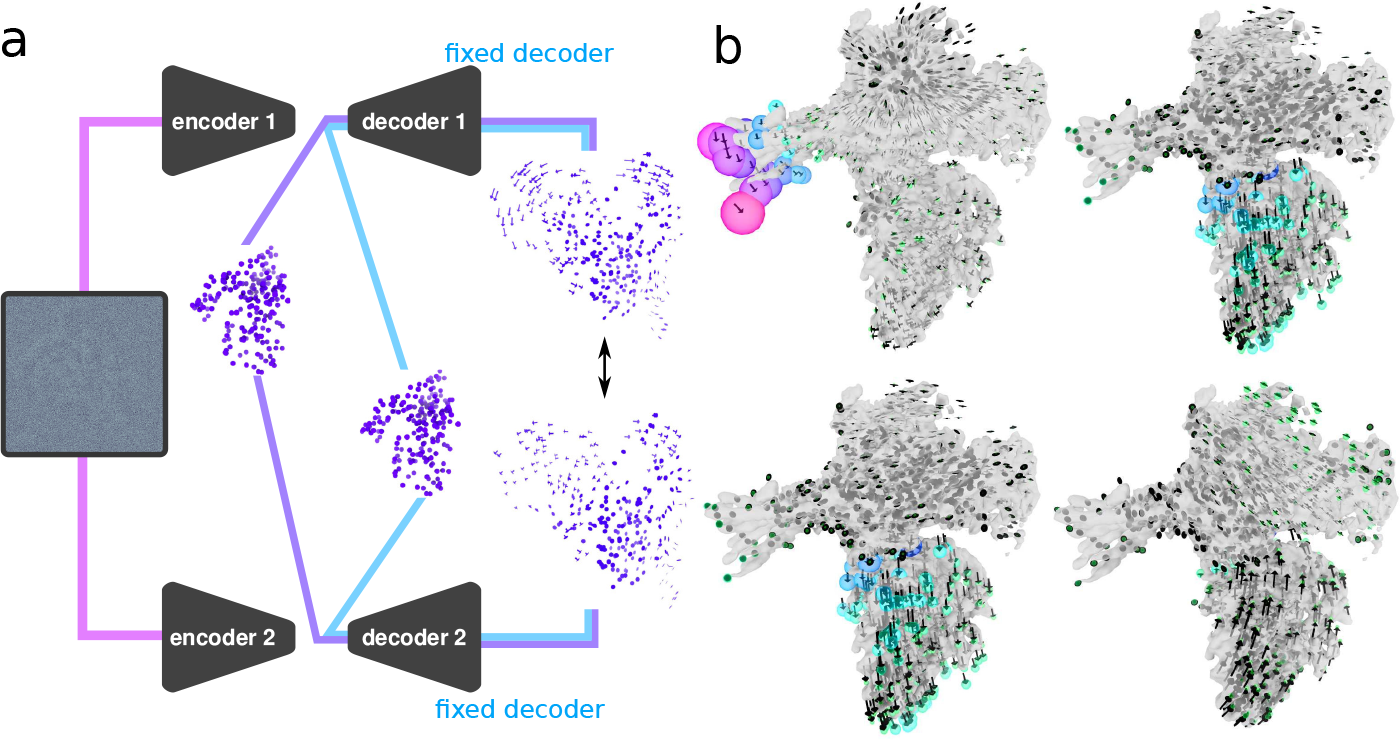
Error estimation for the deformations. **a:** Particles of a validation subset (here 10% of the particles) are fed into both encoders. The encoders are updated, whereas these images are not used for training the decoder. At evaluation time both decoders can be evaluated for the consensus models (purple for the consensus model of half set 1 and blue for the consensus model of half set 2). The resulting displacements can be compared. **b:** Example deformation fields for four particles. The radius of the sphere (coloured by size from blue to pink) at the end of the deformations (black arrows) is determined by the norm of the difference of the deformations from the two decoders.

Besides estimation of deformations, DynaMight also implements a new reconstruction algorithm that aims to correct for the deformations through the reconstruction of an improved consensus map. Reconstruction via (7) only gives an approximation of the minimizer of the convex problem (6). Although it is therefore not guaranteed to yield a useful solution, in practice we observe that DynaMight results in maps with improved local resolutions compared to the standard RELION reconstruction algorithm that assumes structural homogeneity. Nevertheless, our observations that multi-body refinement yields better local resolutions for the complete inner kinetochore complex suggest that there is still room for improvement. It is possible that iterative real-space methods like stochastic gradient descent or algebraic reconstruction techniques, as implemented in Flex3D [12] or Zernike 3D [13], may yield better results. But the iterative approaches would be even more computationally expensive than our weighted backprojection approach, as they may require multiple sweeps through the data and often need optimisation of hyperparameters like the step size. Alternatively, the results with multibody refinement suggest that it may be possible to divide each particle into many smaller “bodies”, and to insert Fourier slices of each of these bodies using orientations that are a combination of the consensus orientation and the average deformation field at that region.

Although many possibilities for improvements still exist, we believe that the current implementation of DynaMight will already be useful. Unlike multi-body refinement, there is no clear minimum of size of moving parts in DynaMight and there is no need for the design of masks that delineate the bodies. In fact, analysis of deformations estimated by DynaMight may assist users to define those masks for subsequent multi-body refinements. The implementation inside RELION-5 will make DynaMight easily accessible to many users, and its wider application will provide feedback for future developments of even better tools to analyse molecular motions in biological macromolecules. The unresolved challenges of how to exploit more prior knowledge, while preventing the pitfalls of model bias, as explored in this paper, imply that this topic will remain an active area of research.

## Acknowledgements

We thank Jake Grimmett, Toby Darling and Ivan Clayson for help with high-performance computing; Carlos Esteve Yagüe, Willem Dieperveen, Carola Schönlieb, Ozan Öktem and Kiarash Jamali for helpful discussions; and David Barford for critical reading of the manuscript. This work was supported by the Medical Research Council (MRC), as part of United Kingdom Research and Innovation (UKRI) (MC UP A025 1013 to S.H.W.S.); and by Wave 1 of The UKRI Strategic Priorities Fund under the EPSRC Grant EP/W006022/1, particularly the “AI for Science” theme within that grant & The Alan Turing Institute. The contribution of T.D. was funded by Cancer Research UK (C576/A14109) and UKRI (MC UP 1201/6) to David Barford. For the purpose of open access, the MRC Laboratory of Molecular Biology has applied a CC BY public copyright licence to any Author Accepted Manuscript version arising.

## Author contributions

J.S. designed and implemented Dynamight, ran all experiments and analysed results; D.K. helped with the design and implementation of Dynamight; A.B. provided help with Python and Napari; T.D. and D.B. provided and analysed yeast kinetochore data; S.H.W.S. provided help with RELION and supervised the project. All authors contributed to writing of the manuscript.

## Code availability

DynaMight is distributed for free under a BSD license and can be downloaded from https://github.com/3dem/DynaMight. It is installed automatically with RELION-5.

